## Supplementary Figures for "Two-stage electro-mechanical coupling of a K_V_ channel in voltage-dependent activation"

**This PDF File Includes:**

Supplementary Figs. 1 to 6

**Supplementary Figures**

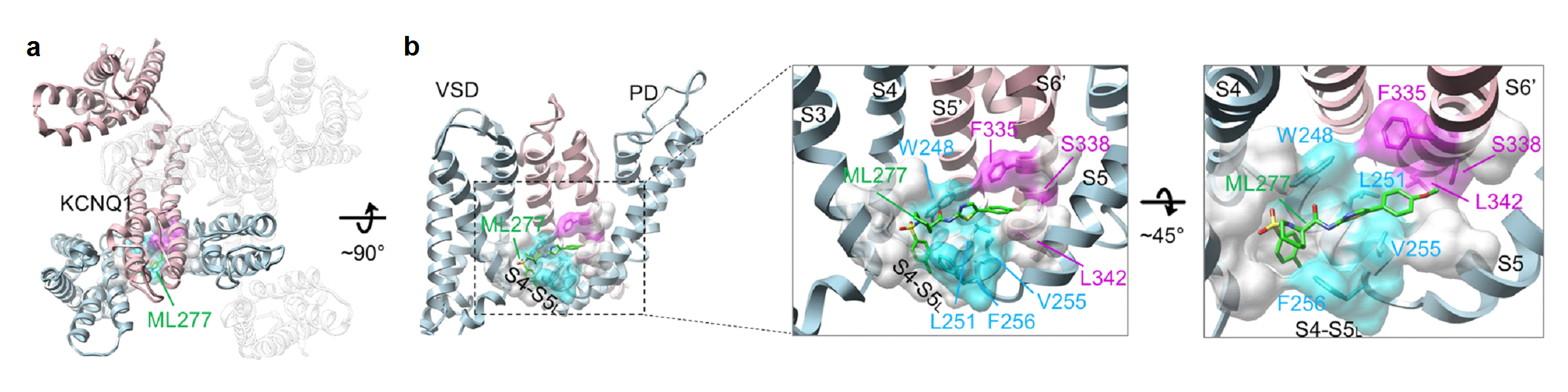

**Supplementary Fig. 1, Molecular docking results of ML277 onto KCNQ1 channel. (a)** Residues interacting with ML277 based on molecular docking and experimental results. Top view of the binding mode of ML277 (green) at the homology model of hK_V_7.1 (see Methods). Two neighboring subunits (light red and light blue) are highlighted. **(b)** Side view of the binding mode of ML277. Only two neighboring subunits (light red and light blue) are shown for clarity. Right panels, detailed structure of the ML277 (green) binding at the S4-S5L/pore interface. Interacting residues that consistent with mutagenesis results shown in cyan (S4-S5L) and magenta (pore).

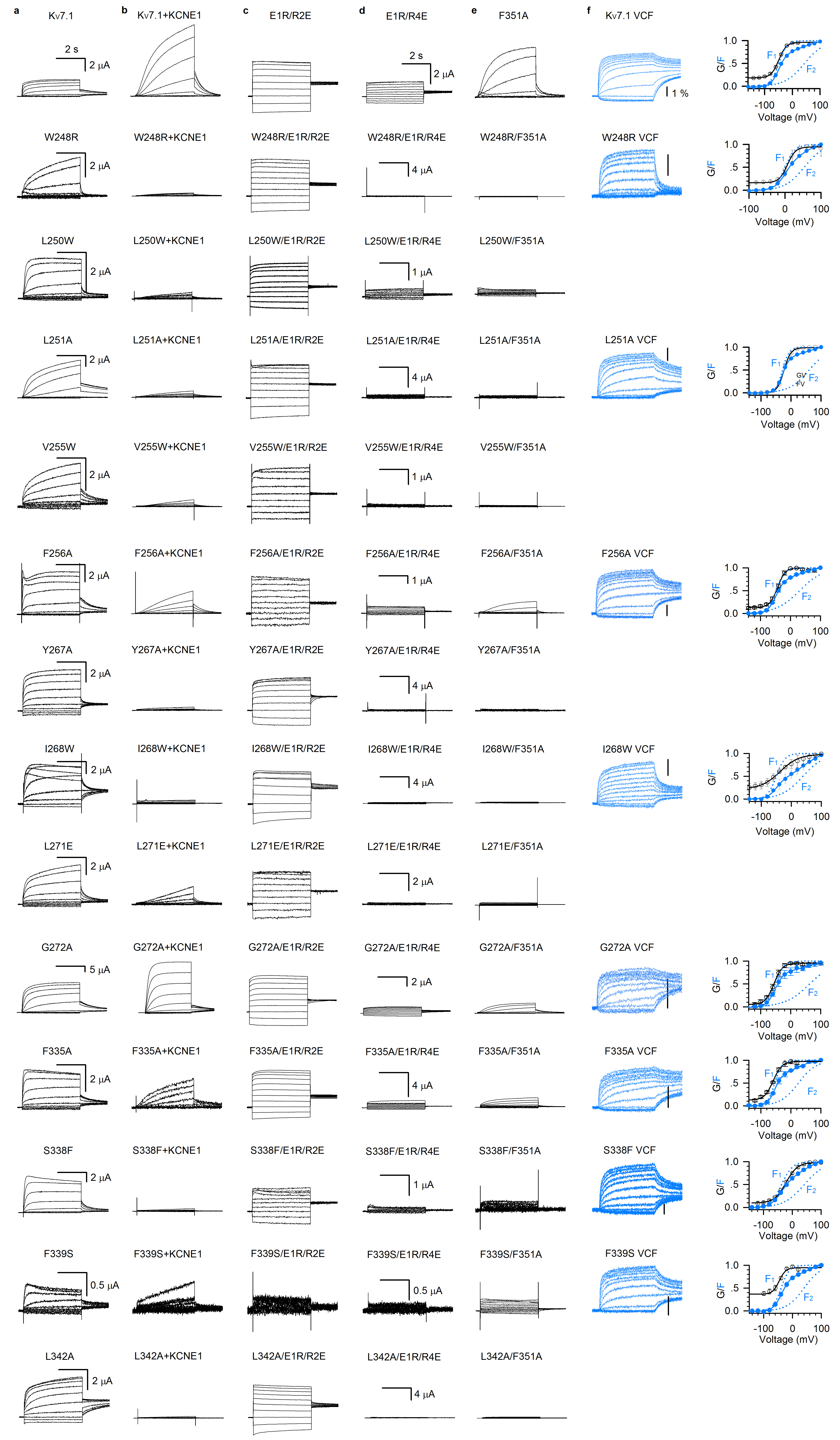

**Supplementary Fig. 2. Dataset for functional validations of AO state E-M coupling residues identified by the ML277 screen. (a-b)** Currents of WT and mutant K_V_7.1 with and without KCNE1. Currents are shown in the same scale. **(c-e)** Currents of WT and mutant K_V_7.1 on top of the IO background E1R/R2E, the AO background E1R/R4E and F351A. Currents are shown in the same scale. **(f)** VCF results of WT and eight mutations in K_V_7.1*. n ≥ 3.

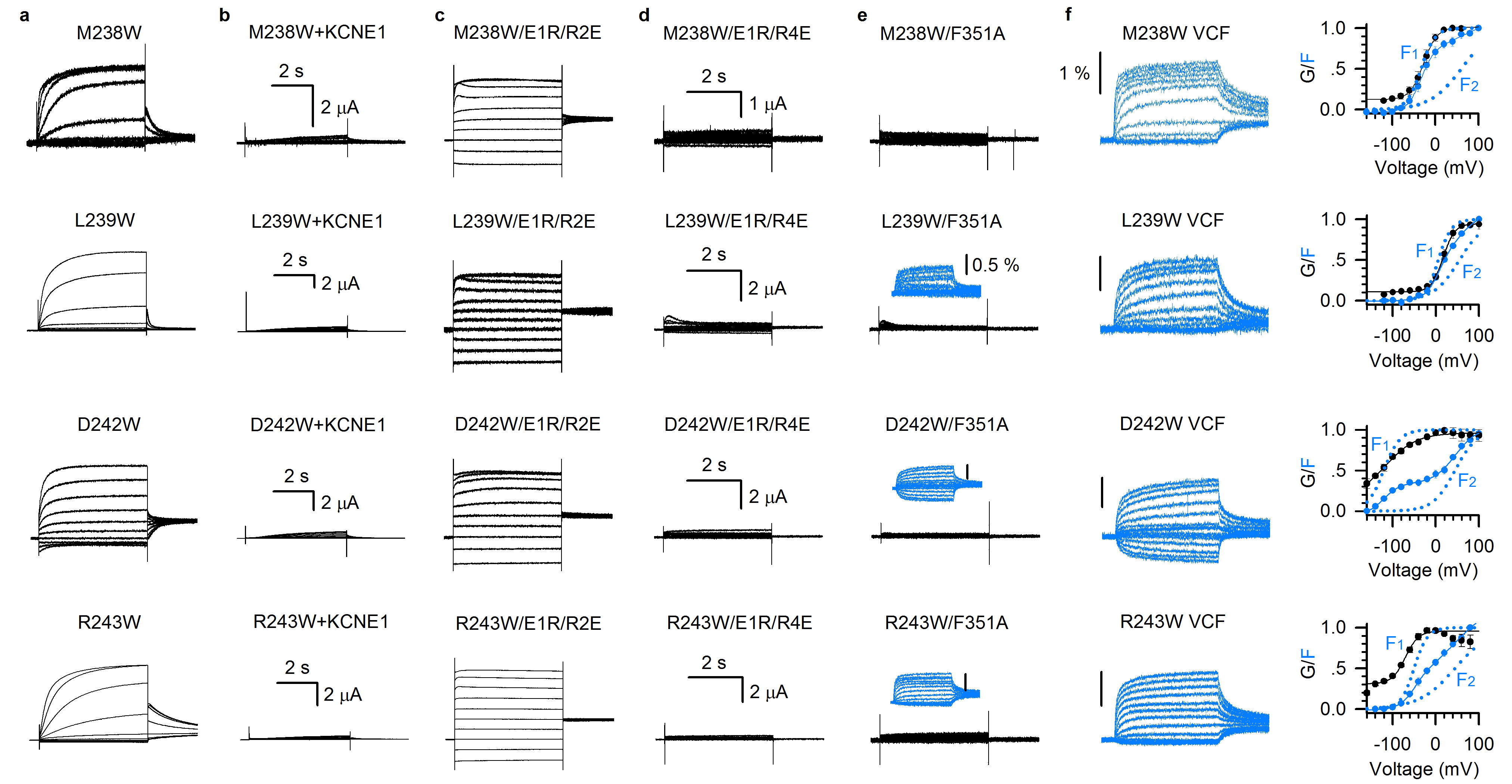

**Supplementary Fig. 3. Dataset for functional validations of S4c residues critical for AO state E-M coupling. (a-b)** Currents of S4c mutations with and without KCNE1. Currents are shown in the same scale. **(c-e)** Currents of the S4c mutations on top of the IO background E1R/R2E, the AO background E1R/R4E and F351A. Currents are shown in the same scale. VCF results (three out of four) are shown in blue. **(f)** VCF results of the four mutations in S4c. n ≥ 3.

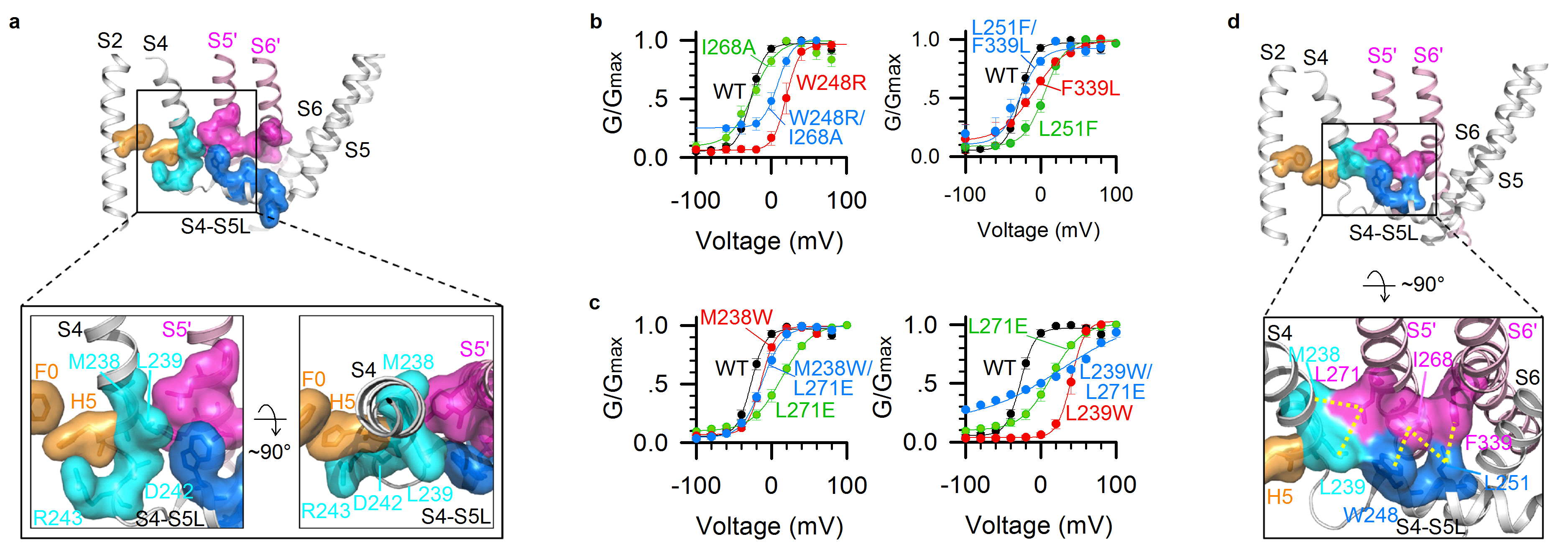

**Supplementary Fig. 4. Double mutant cycle analyses. (a)** Mapping the key residues on to the Kv7.1 cryoEM structure, the VSD of which is at the activated state. Cyan, S4c residues (M238, L239, D242, and R243); Orange, F0 and H5 in the charge transfer center; Blue, S4-S5L residues; Pink, pore residues. Lower panels, detailed structure of the key residues. **(b-c)** Double mutant cycle analysis of the interactions at S4-S5L/pore interface: W248 and I268 (∆∆G= 1.4 kcal/mol) and L251 and F339 (∆∆G= 2.7 kcal/mol), as well as the S4c/S5 interface: M238 and L271 (∆∆G= 1.7 kcal/mol) and L239 and L271 (∆∆G= 4.4 kcal/mol). G–V relations from WT and mutations are fitted with Boltzmann equation. n ≥ 3. **(d)** Mapping the interactions M238/L271 and L239/L271 at the S4c/S5 interface, and W248/I268, L251/I268, and L251/F339 at the S4-S5L/pore interface. Color code is the same as in a, and yellow dashed lines indicate the interactions.

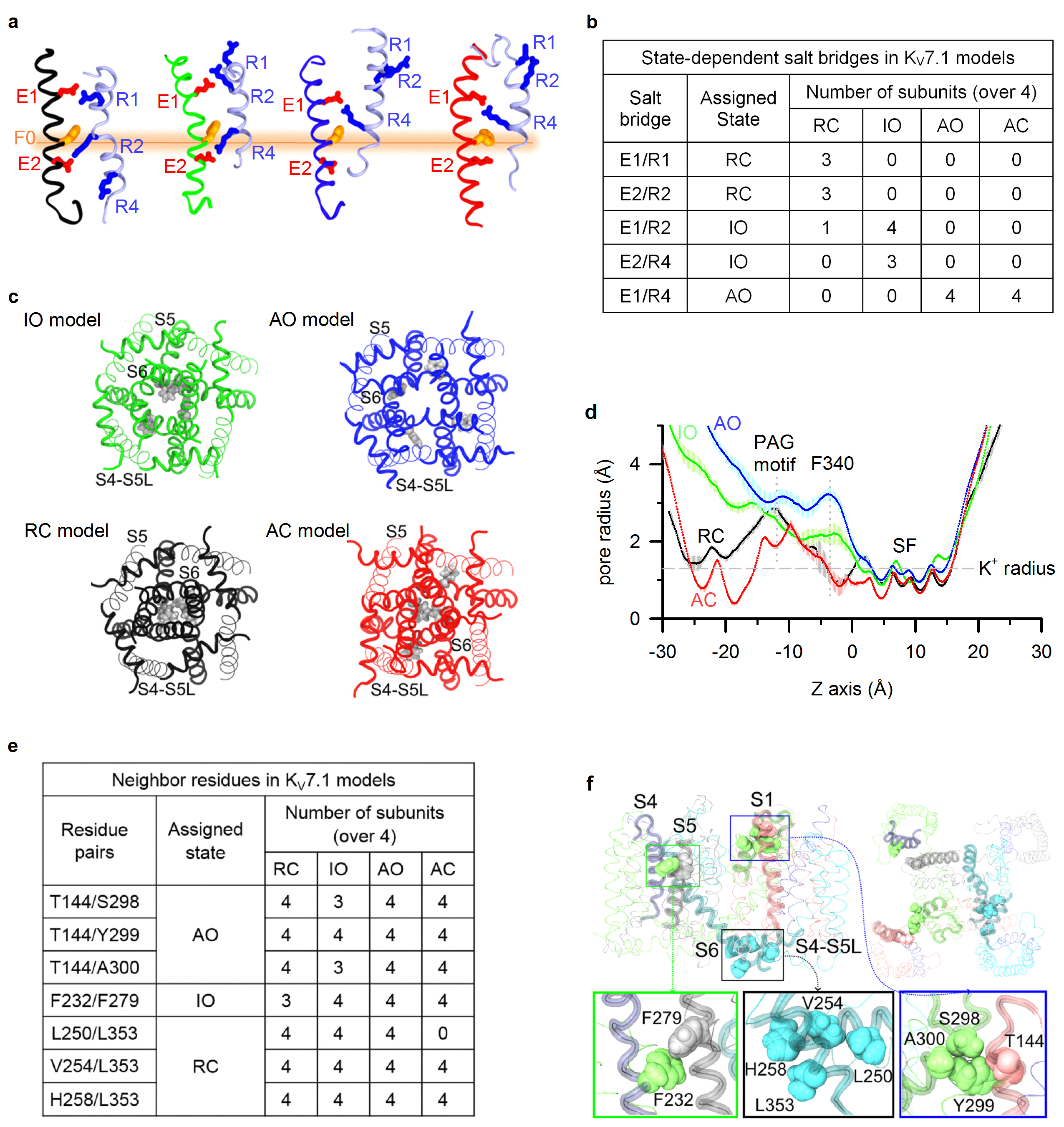

**Supplementary Fig. 5. Structural validation of the MD trajectories obtained for K_V_7.1 models.** **(a)** State-dependent K_V_7.1 VSD salt-bridges^1-4^. Representation of S2 (in black, green, blue and red) and S4 (in blue) segments in K_V_7.1 RC, IO, AO and AC models, respectively. S4 gating charges are represented in blue sticks, while its S2 countercharges are represented in red sticks. The hydrophobic plug F167 (F0) is represented in orange sticks. **(b)** Summary table of state-dependent K_V_7.1 VSD salt-bridges in the subunits of K_V_7.1 models. **(c)** Intracellular view of S4-S5L and PD segments in K_V_7.1 IO (top, left), AO (top, right), RC (bottom, left) and AC (bottom, right) models. The gray spheres are sidechains of F340. **(d)** Average pore radii of the conduction pathways of K_V_7.1 models. Standard deviations are represented by black, green, blue and red horizontal bars for the MD trajectories obtained from RC, IO, AO and AC models, respectively. **(e)** Summary table of state-dependent KCNQ1 neighbor residues pairs identified by cysteine scanning studies and previous double mutant cycle analysis in the subunits of KCNQ1 RC, IO, AO and AC models involving T144 ^5^, F232 ^6^, and L353 ^7^. **(f)** State-dependent pairs of KCNQ1 neighbor residues. In the upper panel, the side view (left) and the extracellular view (right) of the seven pairs of neighbor residues in the KCNQ1 IO model are shown. The transmembrane segments (in cartoon) bearing the residues of interest are highlighted in transparent colors. In the lower panel, the zoomed views of IO state specific interaction in IO model (left), RC state specific interactions in RC model (middle), and of AO state specific residue pairs in AO model (right) are displayed.

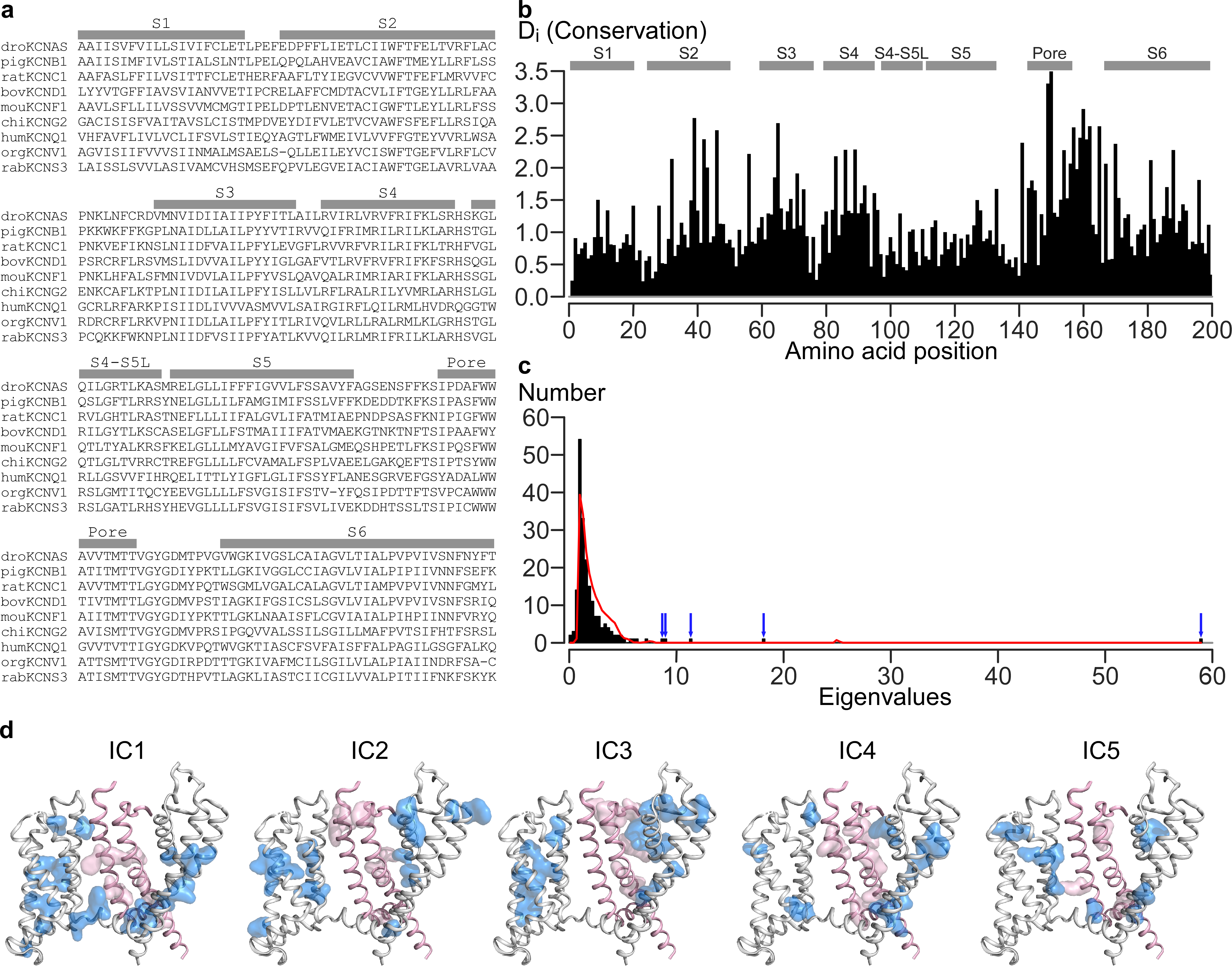

**Supplementary Fig. 6. Statistical coupling analysis of domain-swapped K_V_ channels. (a)** Expanded sample MSA (from Figure 4a) used as input for analysis by SCA. MSA uses the nomenclature established by the HUGO Gene Nomenclature Committee (HGNC) for voltage-gated potassium channels^8^. K_V_ channels can adopt two types of architectures. In “domain-swapped” channels, the VSD from one subunit contacts the PD from the neighboring subunit. In “non-domain-swapped” channels, the VSD from one subunit contacts the PD from the same subunit. These distinct architectures likely give rise to different modes of E-M coupling. SCA was applied to the K_V_ channel superfamily in prior studies^9^; however, the prior studies utilized both domain-swapped and non-domain swapped K_V_ channels in the input data. In this MSA, only domain-swapped K_V_ channels were included. One member from each of K_V_1 – K_V_9 is shown. The final processed MSA includes 1421 sequences at 200 positions. **(b)** First order conservation by amino acid position within the MSA. Bars indicate positions of the helical segments. **(c)** A histogram of eigenmodes tallied by their associated eigenvalues from SCA calculation (black). Red indicates the spectrum calculated by randomizing the input. Five significant eigenmodes can be seen by the black bars rising above the randomized red curve (blue arrows) at high eigenvalues. **(d)** Amino acid residues within independent component 1 through independent component 5 identified by SCA mapped onto the K_V_7.1 cryoEM structure. Blue surfaces indicate all IC positions on one subunit, pink surfaces indicate the same IC positions on the neighboring subunit. Only the S5 and S6 helices are shown on the neighboring subunit for sake of clarity (pink helices). Accordingly, only the IC positions on the neighboring subunit on S5 and S6 are shown (pink surfaces). Using the IC submatrix and based on the strength of external correlation (see Methods, Figure 4c), IC2+IC3 are combined to form sector 1 (Fig. 4d), while IC1+IC4+IC5 are combined to form sector 2 (Fig. 4e).
